# 3D Tissue elongation via ECM stiffness-cued junctional remodeling

**DOI:** 10.1101/384958

**Authors:** Dong-Yuan Chen, Justin Crest, Sebastian J. Streichan, David Bilder

## Abstract

Organs are sculpted by extracellular as well as cell-intrinsic forces, but how collective cell dynamics are orchestrated in response to microenvironmental cues is poorly understood. Here we apply advanced image analysis to reveal ECM-responsive cell behaviors that drive elongation of the Drosophila follicle, a model 3D system in which basement membrane stiffness instructs tissue morphogenesis. Through *in toto* morphometric analyses of WT and ‘round egg’ mutants, we find that neither changes in average cell shape nor oriented cell division are required for appropriate organ shape. Instead, a major element is a reorientation of elongated cells at the follicle anterior. Polarized reorientation is regulated by mechanical cues from the basement membrane, which are transduced by the Src tyrosine kinase to alter junctional E-cadherin trafficking. This mechanosensitive cellular behavior represents a conserved mechanism that can elongate ‘edgeless’ tubular epithelia in a process distinct from those that elongate bounded, planar epithelia.

## INTRODUCTION

A long-time goal of biology is to understand the full set of mechanisms that shape a functional organ. Many morphogenesis studies have focused on only a part of the organ, either by culturing dissected portions *ex vivo*, or by restricting *in vivo* imaging to an optically convenient region. Fundamental morphogenetic principles have emerged from classical examples such as Keller explants of Xenopus embryos ^1^, as well as contemporary examples such as the Drosophila germband ^2^. However, these studies remain limited because creation of artificial boundaries prevents evaluation of outside influences including tissue-wide mechanics. Only now are comprehensive analyses of systems like the Drosophila notum and wing disc, zebrafish gastrula and avian embryo in play ^3^. Nevertheless, these tissues tend to be treated primarily as two-dimensional sheets, in contrast to the many *in vivo* organs that contain multiple tissue types organized in three dimensions (3D). Thus, there is a need to study true 3D organs with *in toto* approaches.

The Drosophila egg chamber, or follicle, provides an excellent model for this goal. Follicles have an architecture that is typical of a number of animal organs, with several components that associate to form a 3D acinar epithelium surrounding a lumen ^4^. At the same time, the simplicity and highly regular development of the follicle lend themselves to comprehensive analyses. The follicle exhibits straightforward and symmetric geometry for much of its development, while its cells originate from only two stem cell populations and show limited differential fates ^5^. Follicles can be genetically manipulated using the powerful Drosophila toolkit, and are well-suited for imaging either in fixed preparations or when cultured live *ex vivo*.

Development of the follicle involves several conserved morphogenetic behaviors including initial primordial assembly, epithelial diversification, and collective cell migration. A major focus for mechanistic studies has been follicle elongation, during which the initially spherical organ transforms into a more tube-like ellipsoid shape ^5,6^. ~2-fold elongation is seen in ~40 hours between follicle budding at st. 3 to the end of st. 8; eventually there is ~2.5-fold overall elongation when the egg is laid ~25 hours later. This degree of elongation is similar to that in paradigmatic morphogenetic systems such as the amphibian neural plate and mesoderm, or the Drosophila germband. In the latter tissues, the main cellular behavior that drives elongation is convergent extension, as cells intercalate mediolaterally toward a specific landmark that is defined anatomically and/or molecularly. However, these tissues have defined borders, which create boundary conditions to instruct and orient cell behaviors. No such boundary is evident along the ‘edgeless epithelium’ of the Drosophila follicle ^7^, and the cellular changes that drive elongation of this acinar organ are not known.

We recently showed that mechanical heterogeneity patterned not within the cells of the follicle, but instead within its underlying basement membrane (BM), instructs organ shape ^8^. Specifically, a gradient of matrix stiffness that is low at the poles and peaks in the organ center provides differential resistance to luminal expansion, leading to tissue elongation. Construction of this pattern relies in part on a collective migration of cells around the follicle equatorial axis, leading to global tissue rotation ^9^. But how the cells of the epithelium respond to stiffness cues and engage in the dynamics that actually elongate the organ along the AP axis remains unexplored. Here we identify an unexpected cell behavior that drives follicle elongation and demonstrate its regulation by BM stiffness cues, thus connecting extracellular mechanical properties to intracellular signaling that drives intercellular morphogenesis *in vivo*.

## RESULTS

### *In toto* morphometries of Drosophila follicles

We established an imaging and computational platform to acquire morphometric data from follicles during their major elongation phase, from st. 4 to st. 8, prior to major asymmetries between the anterior and posterior hemispheres (see **Methods**). Because morphometric measurements can be altered by mounting preparation (e.g. flattened on a slide and/or coverslip) and by artifacts of imaging plane (e.g. XY slices of cells that also have Z orientation in the epithelial plane), complete XYZ images of follicles within a depression slide were captured on a confocal microscope, and analyzed using advanced software including the recently described package ImSAnE ^7,10^ (**Fig. 1a-c**). lmSAnE detects the surfaces of 3D objects and projects them onto 2D planes to facilitate quantitative analysis. Cells were segmented via basal profiles, and projections can be selected that preserve the size or orientation of the epithelial cells with respect to follicle axes (**Fig. 1d-f**; see **Fig. 1g** for definition of meridional and latitudinal axes). The veracity of ImSAnE projections was confirmed by the ability to reconstruct 3D follicles *‘in silico’* (**Fig. 1g and Video 1**). Importantly, this approach allowed complete analysis of the epithelium, including both polar regions, which are seldom scrutinized due to their high curvature and their position perpendicular to the standard imaging plane.

**Fig. 1.**
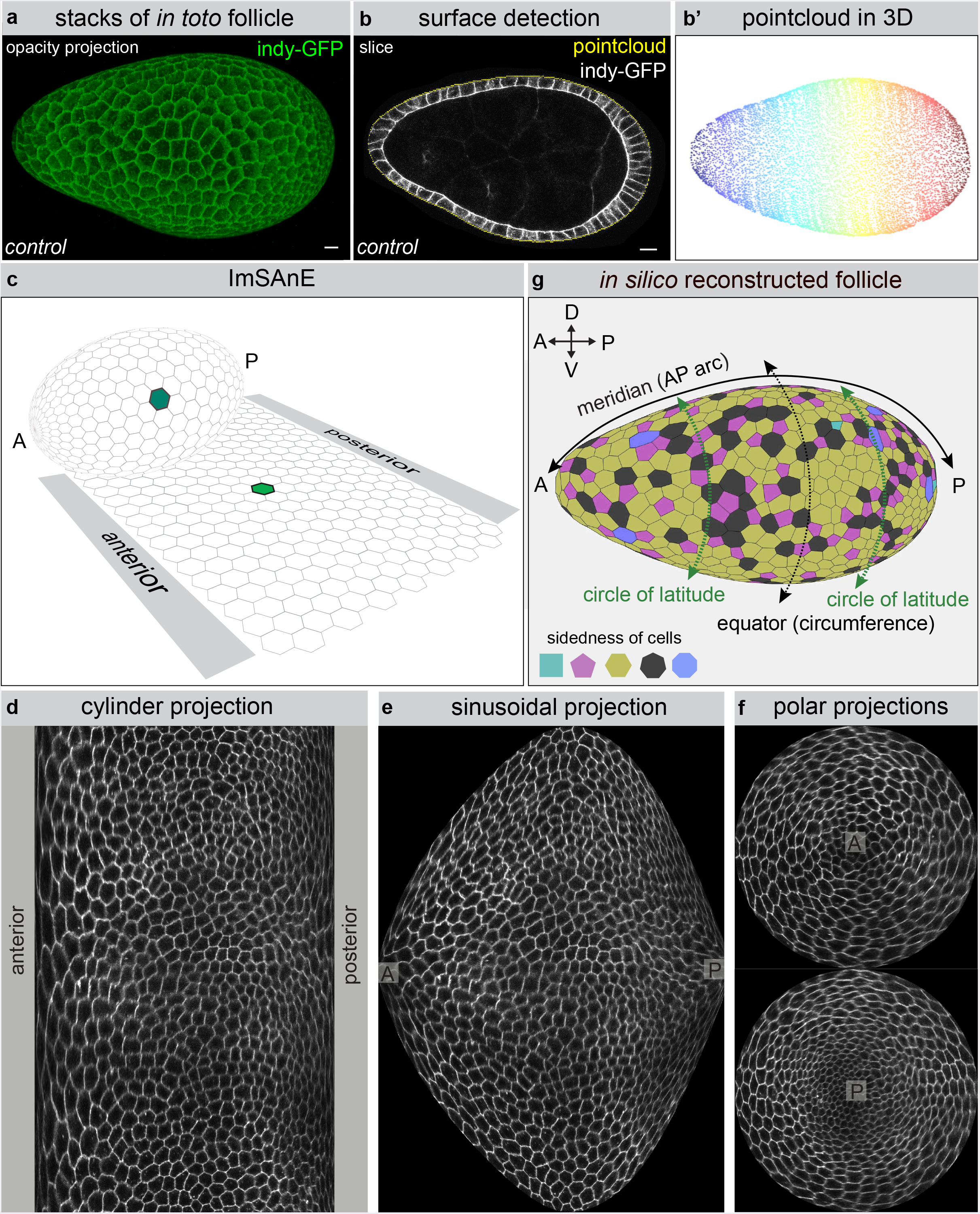
Workflow for follicle morphometrics. **a**, Full 3D stacks of non-compressed Indy-GFP (green) expressing follicles are obtained with confocal microscopy **b**, after which a 3D pointcloud is extracted from the detected surface. **c-f**, ImSAnE then converts the pointcloud into various 2D projections of segmented cells. **g**, Projections and segmentation veracity are confirmed by 3D reconstruction of the follicle (see also **Video 1**). This workflow enables analyses of poles as well as axes along the follicle epithelium, as defined in **g**, meridians connecting anterior (A) and posterior (P) poles, and circles of latitudes including the equator (circumference). Scale bars, 10 μm.

Accurate morphometric comparisons are dependent upon objective identification of developmental stages. We merged classical morphological staging ^11^ with quantitative data obtained above, and generally found good accordance that allowed boundaries based on follicle cell number to be set (**Supplementary Fig. 1**). Stage 6, which is critical for elongation analyses, does not have clearly defined morphological criteria. We subdivided it into stages 6A and 6B to better capture the range of cell numbers and an inferred longer duration, as well as the accelerated rate of elongation in the latter stage (**Fig. 2a, e; Methods**). We also limited our analysis of st. 8 follicles to the presumed youngest half, excluding the larger follicles in which the transition to squamous epithelial cells at the anterior is clearly evident.

**Fig. 2.**
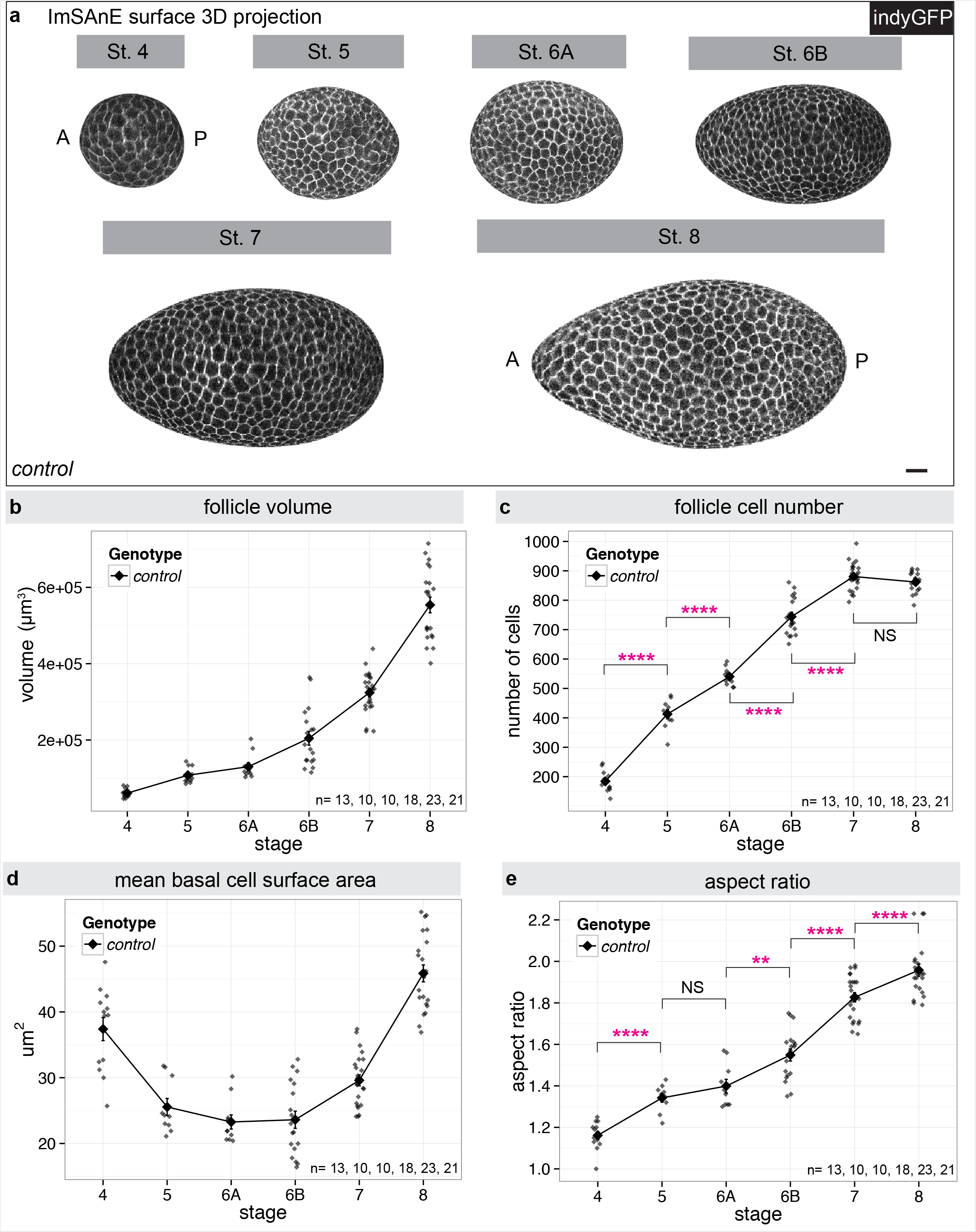
Staging of WT follicles. **a**, Representative ImSAnE surface 3D projection of Indy-GFP (gray) expressing follicles at st. 4-8. Note the increased elongation of st. 6B follicles as compared to st. 6A. Scale bar, 10 μm. **b**, Follicle volume increases ~exponentially during these stages, ~9 fold from st.4-8. **c**, Follicle cell number increases ~linearly with stage, correlates with most morphological criteria, and can be used to define boundaries for each stage. Follicles at stage 7 have reached their maximum cell number. P values= 2.1e-09, 5.1e-06, 1.8e-11, 1.9e-08, and 0.12; two-sided Welch’s t-test. **d**, Mean basal surface area of each follicle cell increases at st. 7 with the transition from cell division to endoreplication. **e**, Kinetics of follicle elongation. P values = 1.9e-06, 0.16, 0.0024, 7.8e-09, and 9.8-e03; two-sided Welch’s t-test. n, Sample size. Error bars, s.e.m. NS, not significant, **P<0.01, ****P<0.0001.

This analysis confirmed, quantified, and refined parameters of WT oogenesis previously derived using other methods. When considered by stage, follicle volume grows roughly exponentially (**Fig. 2b**), while cell numbers in the follicle epithelium increase roughly linearly (**Fig. 2c**), finally reaching 862 ± 32 standard deviation (s.d). cells, similar to both early and more recent estimates ^12,13^. As expected, basal surface area of epithelial cells decreased during division stages and increased after the onset of endoreplication at st. 7 (**Fig. 2d**). Both basal surface areas and apicobasal profiles were similar along the AP meridian through st. 6, suggesting that cells have largely equal volumes. However, at st. 7 basal surface area significantly increased in cells at the anterior one-third and apicobasal profiles decreased, suggesting the first onset of the transition of this population to eventually become fully squamous. Overall, we measure a roughly linear increase in elongation, with an aspect ratio increase from 1.16 at st. 4 to 1.96 at early st. 8, reflecting 54% of the total elongation eventually seen in the mature egg (**Fig. 2e**).

### Cellular behaviors during tissue elongation

We then considered the morphogenetic behaviors that could drive follicle elongation. In other systems, tissue elongation is known to result from oriented cell division, polarized cell shape changes, or oriented cell rearrangements ^13^. We analyzed follicles using ImSAnE to capture parameters for every cell, eliminating artifacts associated with the imaging plane and preserving the orientation along the arcs of the curved epithelium, including at the poles. To analyze the orientation of cell division (**Fig. 3a**), we expressed the microtubule-binding protein Jupiter-GFP to mark the mitotic spindles and midbodies (**Fig. 3b-c and Video 2**); we then live-imaged follicles from st. 3 through st. 6, when mitotic divisions terminate. Midbody position, reflecting the ultimate plane of cytokinesis, was variably aligned in the epithelial plane during st. 3-4, but during st. 5-6 became preferentially aligned along the meridian (**Fig. 3d**). Thus, oriented cell divisions could potentially contribute to follicle elongation prior to st. 7.

**Fig. 3.**
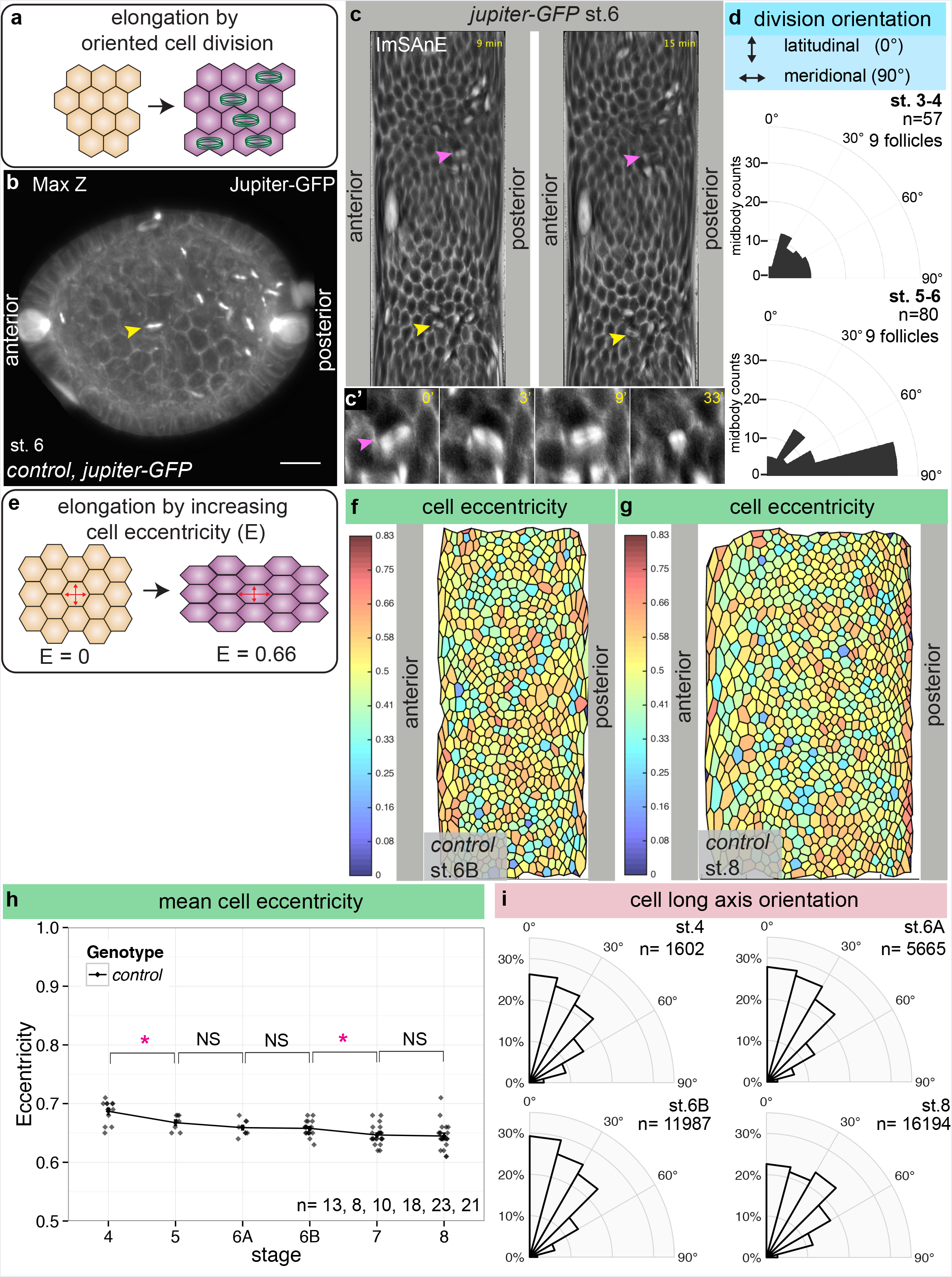
Oriented cell division and cell eccentricity in elongating follicles. **a**, Schematic of tissue elongation via polarized cell division. **b**, Maximum Z projection from confocal stacks and **c**, ImSAnE cylinder projections of the same live-imaged follicle expressing Jupiter-GFP (gray) to mark mitotic spindles and midbodies. Yellow arrowheads mark the same cell division seen in **b** and **c**. Magenta arrowhead in **c** and **c’** marks a cell division in which the spindle changes orientation (see also **Supplementary Fig. 2e**). **d**, Quantitation of division orientation (0°= latitudinal; 90°= meridional) shows that cytokinetic angles are broadly distributed in st. 3-4, round follicles but become oriented along the AP arc in elongating st. 5-6 follicles. e, Schematic of tissue elongation via increased cell eccentricity. **f-g**, ImSAnE projections of st. 6B and st. 8 follicles, colour-coded for eccentricity (0= rounded; 0.83= highly elongated). **h**, Mean cell eccentricity changes modestly during st. 4-8, as compared to the strongly increased elongation of the organ overall. P values = 0.01, 0.17, 0.8, 0.01, and 0.77; two-sided Welch’s t-test. i, Orientation of cell eccentricity (0°= latitudinal; 90°= meridional) shows that most cells are elongated perpendicular to the AP axis. n, Sample size. Error bars, s.e.m. NS, not significant, *P<0.5.

To determine if polarized cell shape changes (**Fig. 3e**) account for overall follicle elongation, we measured the eccentricity of each follicle cell; that is, the aspect ratio of its best-fit ellipse (**Fig. 3f-g**). During st. 5-8, even though the distribution of cell eccentricity varies, follicle cells maintain a quite consistent mean eccentricity, between 0.65-0.7 (**Fig. 3h**). Importantly, we also determined the orientation of cell eccentricity within the epithelial plane and found that at all stages, most cells’ long axes are oriented latitudinally (**Fig. 3i**). Cells therefore tend to be elongated perpendicular to, rather than parallel to, the axis of tissue elongation, and their average shapes change only slightly during a near doubling of organ aspect ratio.

To explore whether cell rearrangements (**Fig. 4a**) occur during follicle elongation, we first analyzed cellular topology in fixed follicles (**Fig. 4b**). Changes in topological order can be caused by either cell proliferation or by neighbor exchange, and ordered tissues have a greater proportion of hexagonal cells ^14^. St. 4 follicles were the most disordered, likely reflecting their high rate of cell division, rather than tissue elongation which is limited at this time. Topological order then increased, providing evidence for rearrangement of cell junctions (**Fig. 4c**).

To seek direct evidence of cell rearrangements, we counted cells along the follicle arcs (**Fig. 4d**). These counts revealed a 65% increase in the ratio of meridian to equatorial cells between st. 6-8 (**Fig. 4e**). Moreover, in post-mitotic follicles between st. 7-8, an addition of ~3 epithelial cells along the AP meridian was seen (**Fig. 4f**). These data demonstrate that cell rearrangements do occur during follicle elongation, leading to intercalation of cells from the equatorial to the AP axis.

We next searched for intercalation events by examining follicles live-imaged *ex vivo*. We did not see rosette-like arrangements, and while we saw occasional 4-cell (T2-like) junctions ^15,16^, we were unable to consistently identify T1>T3 transitions in movies from st. 6-8. However, live-imaged follicles at these stages showed signs of deterioration after 6 hours, only ~1/4 of the period in which elongation takes place, limiting our ability to draw conclusions from this approach.

**Fig. 4.**
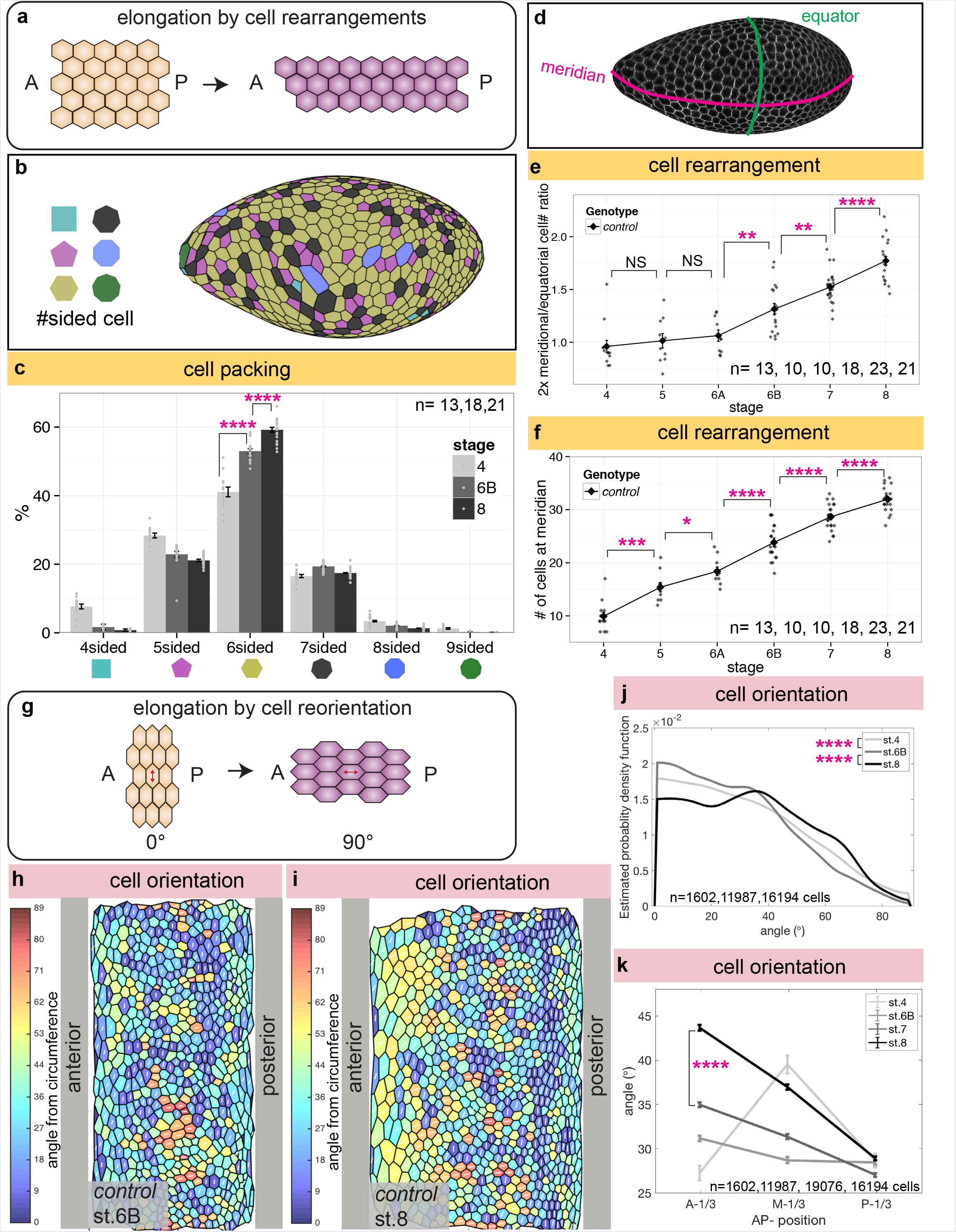
Cell rearrangements and cell reorientation in elongating follicles. **a**, Schematic of tissue elongation via cell rearrangements. **b**, Reconstructed st. 8 follicle colour-coded for cell topology. **c**, Quantitation shows increases in topological order (i.e. frequency of hexagonal cells) from st. 4 to st. 8. P values = 6.5e-07 and 2.8e-07; twosided Welch’s t-test. **d-f**, Cell counts demonstrate increases in the ratio of meridional cells to equatorial cells from st. 6B through st. 8; increases in the number of meridional cells also demonstrate cell intercalation. In **e**, P values = 0.56, 0.58, 0.003, 0.0024, and 2.2e-05; two-sided Welch’s t-test. In **f**, P values = 1.58e-04, 0.02, 7.8e-05, 2.0e-05, and 4.2e-05; two-sided Welch’s t-test. **g**, Schematic of tissue elongation via cell reorientation: anisotropically-shaped cells remodel cell junctions to shift the direction of their long axis. h-i, ImSAnE cylinder projections of st. 6B and st. 8 follicles, colour-coded for cell orientation (0 ° = latitudinal; 90 ° = AP). j, Graph showing distribution of cell orientations in WT follicles: estimated density function (y axis) reflects the relative frequency of cells with a given long axis orientation (x axis, as defined in **g**). From st. 4-8, the proportion of AP-oriented cells increases significantly. P values = 4.7e-06 and 3.27e-83; Two-sample Kolmogorov-Smirnov test. **k**, Cells in the anterior one third of the follicle between st. 6B and 8 reorient to become increasingly AP oriented (A-1/3: anterior one third; M-1/3: middle one third; P-1/3: posterior one third; values are average orientation for each region). P value = 4.22e-86; two-sided Welch’s t-test. n, Sample size. Error bars, s.e.m. NS, not significant, *P<0.5, **P<0.01, ****P<0.0001.

Although the average shapes of follicle cells are persistent, their anisotropic dimensions raise the possibility that changes in orientation could be an alternative mechanism for tissue elongation (**Fig. 4g**). Strikingly, as oogenesis precedes, a population of cells reorient their long axis, from primarily along the circles of latitude to increasingly along the AP meridian (**Fig. 4h-j**). Between st.7 and st. 8, reorientation was particularly evident in the anterior third of the follicle (**Fig. 4k**), where the average angle of orientation towards the meridian increases by 8.7°. This cell reorientation, which is coincident with a period of post-mitotic tissue elongation, is thus a candidate contributor to it.

### Cell behaviors in elongation-defective mutants

To test the functional role of the above behaviors, we analyzed them in mutant backgrounds that altered either the behaviors specifically or follicle elongation in general. The most frequently-used elongation mutant disrupts *fat2*, which encodes an atypical cadherin required for follicle planar cell polarity and tissue rotation ^17,18^. *Fat2* mutant or RNAi-depleted follicles show discontinuously variable stiffness ^8,19^, and their elongation diverges from WT follicles at st. 6B; by st. 8 they have aspect ratios of 1.55± 0.08 s.d. instead of 1.96± 0.13 s.d. (**Fig. 5a-c**). Growth of fat2-depleted follicles and proliferation of their epithelial cells are unchanged compared to WT, with the exception of having slightly more cells at st.8 (**Supplementary Fig. 2a-b**). Since total surface area of *fat2*-depleted follicles remains similar to WT (**Supplementary Fig. 2c**), the increase in cell numbers leads to a slight reduction in mean basal surface area of individual cells (**Supplementary Fig. 2d**).

**Fig. 5.**
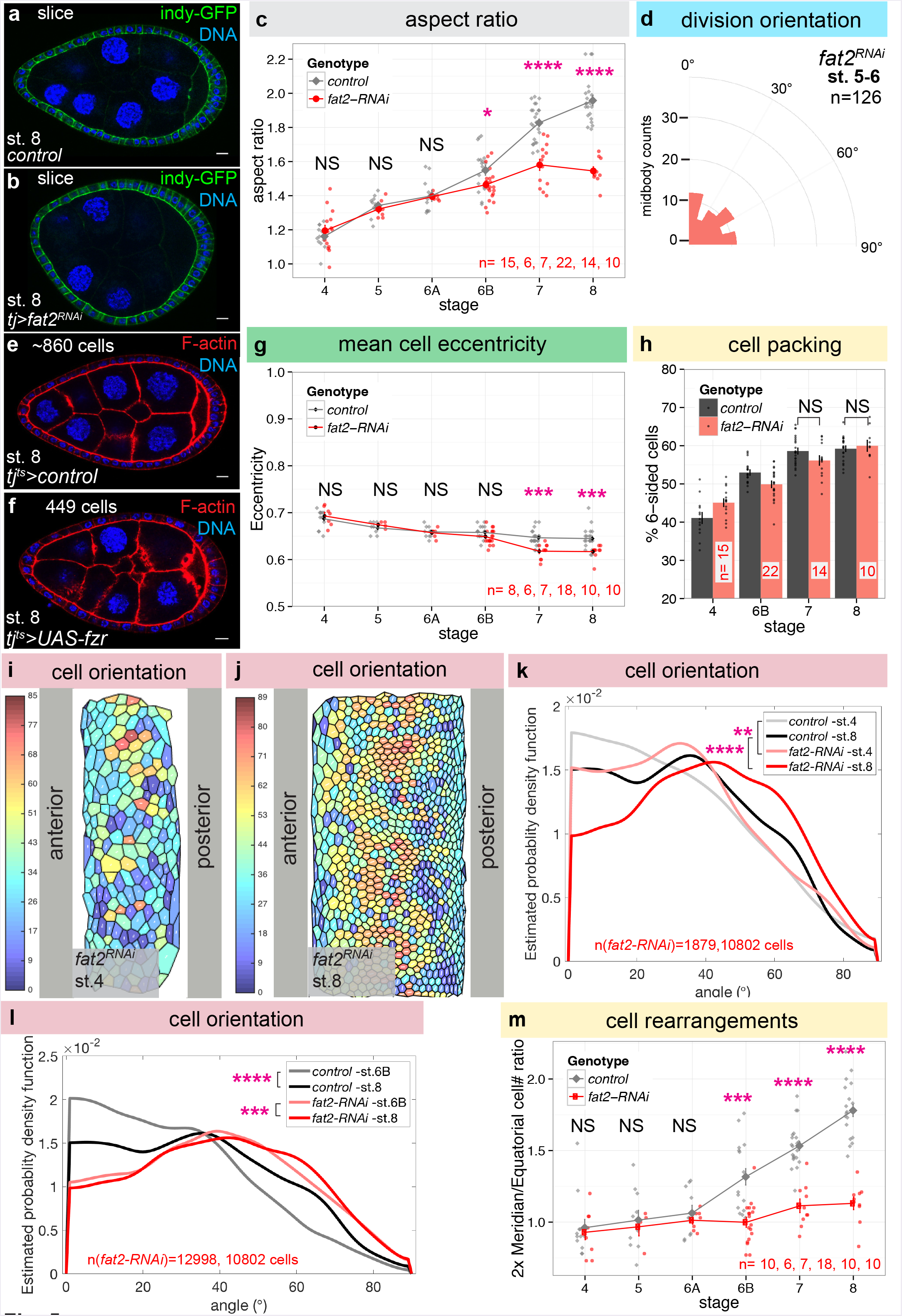
Morphometrics of elongation-deficient follicles. **a-b**, Control(a) and *fat2*-depleted(**b**) follicles at st. 8. **c**, Aspect ratio of fat2-depleted follicles diverges from WT at st. 6B. P values = 0.36, 0.52, 0.89, 0.02, 1.45e-06, and 1.35e-11; two-sided Welch’s t-test. **d**, Cell divisions remain ~randomly oriented in st. 5-6 *fat2*-depleted follicles. e-f, Control (e) and Fzr-overexpressing (f) follicles at st. 8. Premature onset of endoreplication via acute expression of Fzr ~halves follicle cell number, but does not prevent tissue elongation. **g**, Cell eccentricity in *fat2*-depleted follicles does not differ from WT until st. 7, when cells become slightly less elongated. P values = 0.68, 0.67, 0.73, 0.08, 2.2e-04, and 3e-04; two-sided Welch’s t-test. **h**, Topology analysis shows that cells in st. 8 *fat2*-depleted follicles can rearrange to reduce tissue disorder. P values = 0.10, and 0.64; two-sided Welch’s t-test. **i-j**, ImSAnE cylinder projections of st. 6B and st. 8 *fat2*-depleted follicles, colour-coded for cell orientation (0 ° = latitudinal; 90 ° = meridional). **k-l**, Distributions show that *fat2*-depleted follicles show aberrant orientation distributions at st. 4, and do not show the reorientation evident in WT at st. 8. In **k**, P values = 0.0026 and 4.0e-66; Two-sample Kolmogorov-Smirnov test. In l, P values = 3.27e-83 and 6.18e-04; Two-sample Kolmogorov-Smirnov test. m, Cell counts reveal that *fat2*-depleted follicles fail to rearrange cells from circumferential to meridional. P values = 0.56, 0.69, 0.42, 3.28e-05, 1.40e-06, and 1.55e-10; two-sided Welch’s t-test. n, Sample size. Scale bars, 10 μm. Error bars, s.e.m. NS, not significant, *P<0.5, **P<0.01, ***P<0.001, ****P<0.0001.

When analyzing cell divisions in *fat2*-depleted follicles at st. 5-6, we found that, in contrast to WT, the plane of cytokinesis did not orient along the AP arc, but remained randomly aligned as it is in st. 3-4 (**Fig. 5d**). Because fat2-depleted follicles show disrupted BM stiffness at st. 5, before defects in elongation are evident ^8^, we considered the possibility that the oriented divisions seen in st. 5-6 WT follicles are a consequence of the stiffness gradient, rather than a cause of tissue elongation. We therefore examined the initial mitotic spindle orientation in WT follicle cells and found that it was not preferentially AP oriented, even at st. 5-6. Instead, AP alignment of the spindle, often perpendicular to the initial cell long axis, generally occurred after metaphase, consistent with the outcome of division being organized by tissue-wide tension (**Supplementary Fig. 2e and Video 3**). Additionally, RNAi to *mushroom body defective (mud)*, which regulates the mitotic plane in follicle cells ^17^, did not induce elongation defects (st. 8 aspect ratio=1.86 n=3). Finally, we inhibited mitoses in otherwise WT follicles by overexpressing

*fizzy-related (fzr)*, a Cdh1 homolog that is sufficient to switch cells from proliferative to an endocycle program ^18^. Fzr overexpression from st. 4 limited epithelial cell numbers to only 415 ± 51 s.d. cells, which is ~48% of WT. Nevertheless, these follicles elongated similar to WT (st. 8 aspect ratio=1.89 n=5, **Fig. 5e-f**). We conclude that oriented cell divisions are not required for follicle elongation.

We then investigated cell shapes, rearrangements, and orientation changes in *fat2*-depleted follicles. Despite the fact that Fat2 is required for follicle rotation, cell eccentricity was similar to WT from st.4 to st.6B (**Fig. 5g**), demonstrating that elongated cell shapes are not dictated by collective cell migration. Cell topology at st. 7-8 also did not differ from WT (**Fig. 5h**), indicating that *fat2* cells are capable of rearranging to increase tissue order. However, the initial distribution of cell orientations in *fat2* follicles differed from WT even at st. 4-5: cells orient their long axis more along the AP meridian, with a broader distribution of orientations than the highly skewed distribution of WT (**Fig. 5i-k**). This phenotype can be seen before significant differences in organ elongation are evident and represents the earliest morphometric defect seen in *fat2* follicles. Moreover, whereas the distribution of orientations in WT follicles shifts substantially following st. 6B, *fat2-depleted* follicles show only a minor shift (**Fig. 5l**). The shift in WT follicles changes average cell orientation 5 times that seen in fat2-depleted follicles (5° vs 1°). Finally, equatorial and meridional cell counts to assess intercalation during st. 6-8 revealed that the ratio did not increase in *fat2*, as it did in WT (**Fig. 5m**). These data suggest a relationship between changes in cell orientation and the cellular rearrangements that elongate the follicle.

### Rack1 regulates Src to reorient cells in response to BM mechanical cues

The mechanically patterned BM proposed to sculpt the tissue ^8^ is disrupted in the fat2-depleted follicles analyzed above. To uncover how elongation-driving cell behaviors normally respond to these ECM cues, we searched for mutants that are defective in elongation but nevertheless retain graded BM stiffness. We exploited an ongoing genetic screen in the lab in which transgenic RNAi lines are expressed specifically within the follicle epithelium, and those that perturb oogenesis are scored as ‘hits’. One hit targeted *Receptor for Activated Protein Kinase C 1 (Rack1)*, which gave rise to round follicles and round eggs with a high degree of penetrance (**Fig. 6a-d**). Elongation defects emerged at st. 6B, similar to fat2-depleted follicles (**Supplementary Fig. 3a**). This phenotype was seen with two independent RNAi lines targeting different regions of the transcript, and was rescued by overexpression of a Rack1-encoding transgene (**Supplementary Fig. 3b-c**). It was further confirmed by making mitotic clones of *Rack1* null cells in the epithelium, where moderately-sized clones altered follicle aspect ratio (**Supplementary Fig. 3d**). Thus, Rack1 is a new regulator of egg elongation.

**Fig. 6.**
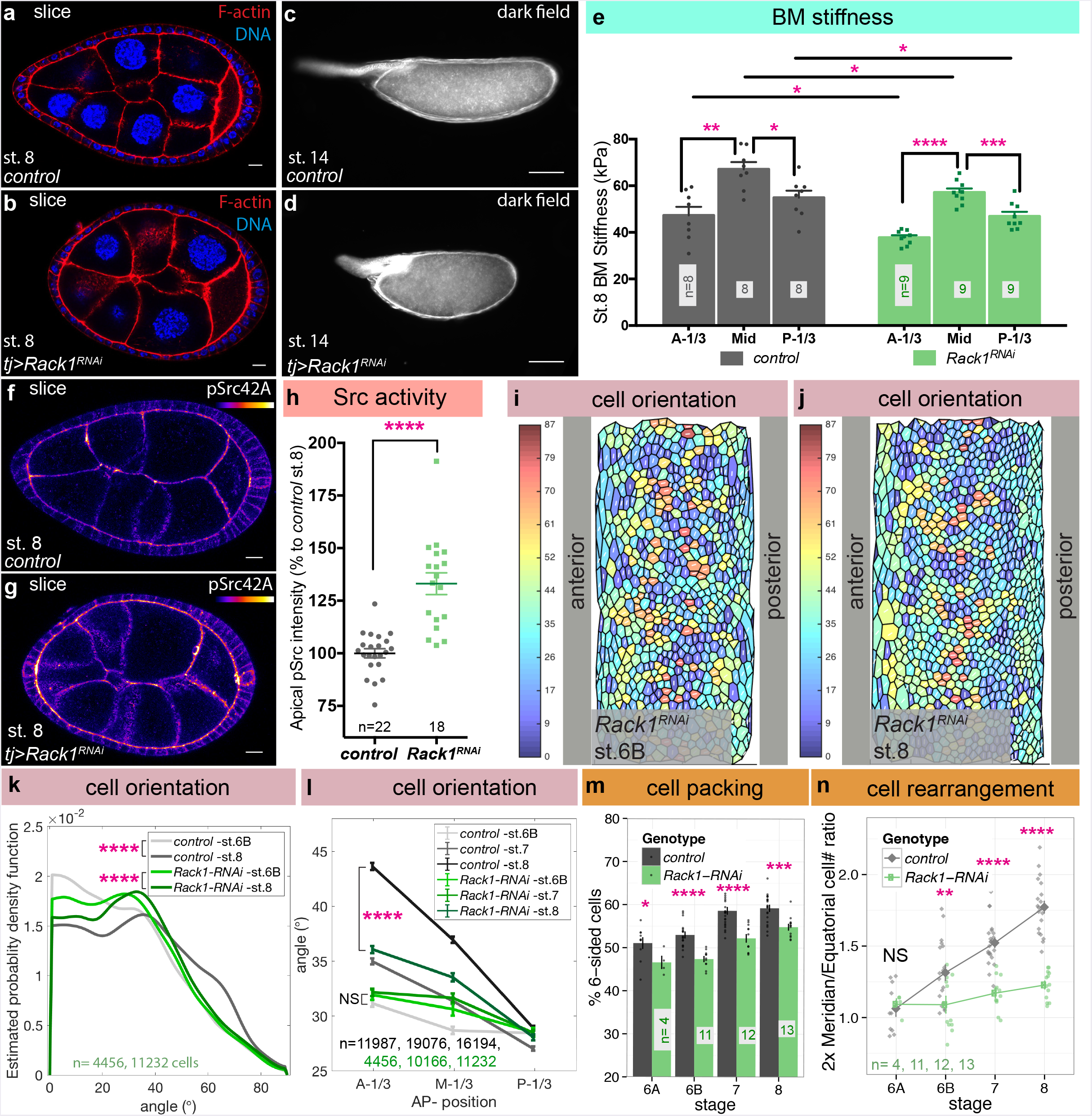
Src regulator Rack1 controls cell reorientation and follicle elongation. **a-d**, RNAi-mediated depletion of *Rack1* produces round follicles (**b**) and round eggs (**d**) compared to controls (**a, c**). **e**, Atomic Force Microscopy (AFM) analysis of the basement membrane of Rack1-depleted follicles shows preservation of an AP gradient of stiffness, although average stiffness is slightly decreased (A-1/3: anterior one third; M-1/3: middle one third; P-1/3: posterior one third). P values between *control* at different positions = 0.0011 and 0.013, between *Rack1^RNAi^* at different positions = <0.0001 and 0.0009, between control and Rack1 ^*RNAi*^ = 0.037(A-1/3), 0.016(M-1/3), and 0.045(P-1/3); two-sided Welch’s t-test. **f-h**, *Rack1*-depleted follicles (**g**) show increased levels of phosphorylated Src (fire LUT) at apical junctions compared to WT (**f**). P values = <0.0001; two-sided Welch’s t-test. **i-k**, ImSAnE cylinder projections of st. 6B and st. 8 *Rack1*-depleted follicles, colour-coded for cell orientation (0 ° = latitudinal; 90 ° =AP), along with distributions. P Value (in **k**) = 3.27e-83 and 8.97e-06; Two-sample Kolmogorov-Smirnov test. l, Cells in the anterior third of Rack1-depleted follicles fail to reorient towards the AP (A-1/3: anterior one third; M-1/3: middle one third; P-1/3: posterior one third). P Values between st.6B *control* and *Rack1^RNAi^* = 0.18 (A-1/3), 0.004 (M-1/3), and 0.91(P-1/3); twosided Welch’s t-test. P Values between st.8 *control* and *Rack1^RNAl^* = 1.51e-58 (A-1/3), 8.31e-13 (M-1/3), and 0.01 (P-1/3); two-sided Welch’s t-test. m, Rack1-depleted follicles impair to rearrange cells to reduce tissue disorder. P Values = 0.046, 1.6e-06, 9.3e-09, and 0.0001; two-sided Welch’s t-test. **n**, Rack1-depleted follicles fail to rearrange cells from the equatorial to latitudinal. P Values = 0.66, 0.00962, 2.67e-08, and 2.10e-12; twosided Welch’s t-test. Error bars, s.e.m. n, Sample size. NS, not significant, *P<0.5, **P<0.01, ***P<0.001, ****P<0.0001. **a-b, f-g**, scale bars, 10 μm and **c-d**, scale bars, 100 μm.

Analysis of *Rack1*-depleted follicles revealed that they did not differ from WT in several assays. Growth of follicles was similar to WT (**Supplementary Fig. 3e-f**). Proliferation of epithelial cells was in general unchanged, except for the increase in st. 8 cell number also seen in elongation-defective *fat2* (**Supplementary Fig. 3g**). In *ex vivo* culture, Rack1-depleted follicles rotated with normal speed and trajectory (**Supplementary Fig. 3h and Video 4**). A reporter for JAK-STAT signaling, which is required for elongation ^8,20^, displayed WT pattern at both poles (**Supplementary Fig. 3i-j**). At st. 6, cell divisions were oriented along the elongation axis, comparable to WT follicles (**Supplementary Fig. 3k**).

We then assayed BM mechanics in Rack1-depleted follicles. Direct Atomic Force Microscopy (AFM) measurements of the BM showed a modest reduction in overall stiffness, but AP anisotropy remained, as in WT (**Fig. 6e**). This contrasts strongly with the disorganized stiffness of *fat2* follicle BMs ^8,19^. When challenged with osmotic pressure induced by placement in distilled water, Rack1-depleted follicles burst slightly more frequently than WT follicles, but not as rapidly as *fat2* follicles ^8^ (**Supplementary Fig. 3l**); bursting position also more closely resembled WT. The overall mechanical properties of Rack1-depleted follicles resemble those seen in follicles overexpressing SPARC or carrying *fat2* hypomorphic alleles that delete only the intracellular domain of the protein, yet these latter genotypes elongate normally, while Rack1-depleted follicles do not ^8,21^. This contrast suggests that Rack1 could be required for cellular behaviors that are triggered in response to the BM stiffness gradient.

Rack1 is a scaffolding protein whose seven WD-40 domains interact with the Src tyrosine kinase, maintaining it in an inactive state ^22^. We asked whether Rack1 might negatively regulate Src activity during follicle elongation. The Drosophila genome encodes two Src orthologs, Src42 which is expressed in follicle cells, and Src64 which is not ^23,24^. In WT follicle epithelia, active phosphorylated Src (pSrc) was found primarily at adherens junctions (AJs) (**Fig. 6f**). AJ staining of pSrc was significantly elevated in Rack1-depleted follicles (**Fig. 6g-h**); total Src42 levels were also somewhat elevated (**Supplementary Fig. 4a-c**). Moreover, overexpression of active Src42 was sufficient to induce elongation defects in otherwise WT follicles (**Supplementary Fig. 4d-e**), while reducing Src42 activity via heterozygosity for a null allele partially rescued the elongation defect in Rack1-depleted follicles (**Supplementary Fig. 4f**).

These results suggest that *Rack1* loss and associated Src activation alters morphogenetic cell behaviors. To identify these behaviors, we analyzed cell and junctional morphology in *Rack1* follicles. While cell eccentricity from st. 6-8 was slightly higher than WT (**Supplementary Fig. 4g**), the strongest defects were seen in cell orientation. At st. 6A and 6B, orientation in Rack1-depeleted follicles was comparable to WT, and did not show the altered distributions seen in *fat2* (**Fig. 6i, k**). However, the reorientation of cells seen during st. 7-8 in WT was notably impaired in Rack1-depleted follicles (**Fig. 6j-k**); in particular, the anterior cells that realign towards the AP axis in WT fail to do so in Rack1-depleted follicles, remaining more latitudinally aligned (**Fig. 6l**). In contrast to WT and *fat2, Rack1* depletion also did not resolve tissue disorder as reflected in cell topology, with lower percentages of hexagonal cells at st. 8 (**Fig. 6m**). Finally, cell counts along the follicle arcs established that Rack1-depleted follicles fail to intercalate cells along the AP meridian (**Fig. 6n**).

Because Rack1 and Src have been implicated in cell junction remodeling in other systems ^20-24^, we analyzed junctional dynamics by performing Fluorescence Recovery After Photobleaching (FRAP) on follicles expressing natively GFP-tagged E-cadherin (Ecad-GFP) ^25^ (**Fig. 7a and Video 5**). While steady-state levels of Ecad-GFP did not differ between WT and Rack1-depleted follicles, FRAP analysis revealed changes in Ecad-GFP dynamics. No differences in the mobile fraction were seen between the two genotypes (**Fig. 7b**), but recovery was significantly slowed in Rack1-depleted follicles, with a 67% increase in the recovery half-time (**Fig. 7c**). Recovery in both genotypes after photobleaching was not due to lateral diffusion (**Supplementary Fig. 4h**). Taken altogether, these data are consistent with a model in which Rack1 regulates Ecad trafficking to permit junctional rearrangements that elongate the tissue.

**Fig. 7.**
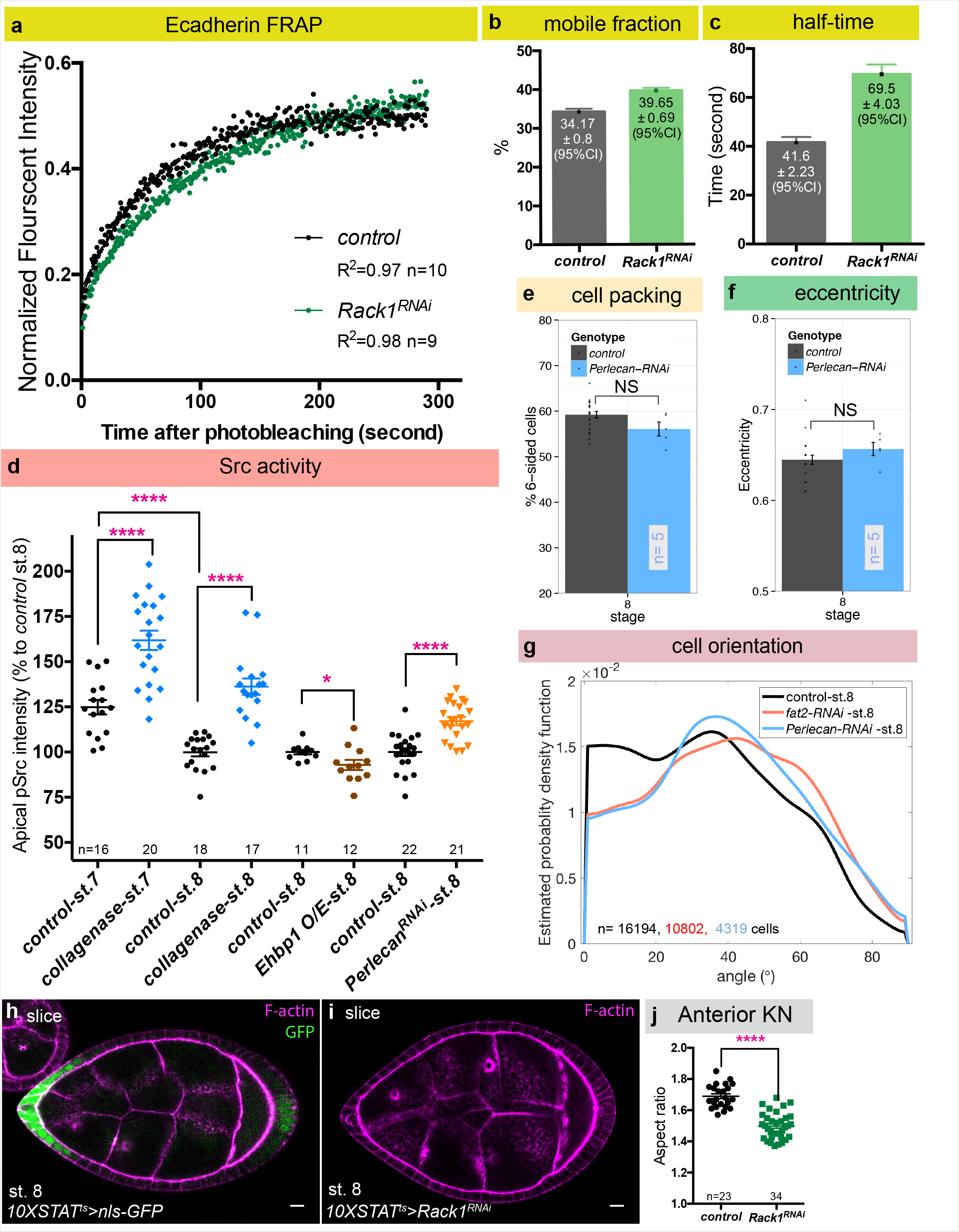
BM stiffness-responsive Src activation increases AJ dynamics. **a**, FRAP analysis of Ecad-GFP in control and Rack1-depleted follicles at st. 8. Dots represent mean of normalized intensity after photobleaching and lines are fitted curves with error bars showing 95% confidence intervals. (see also **Video 5**). **b-c**, Mobile fraction of Ecad-GFP is similar in both genotypes, but recovery half-time is increased when Rack1 is depleted. Error bars, 95% confidence intervals. **d**, Quantitation of pSrc levels shows that reduced BM stiffness is associated with increased Src activity (control st. 7 versus st. 8, collagenase treatment, and depletion of Perlecan). All P values <0.0001; two-sided Welch’s t-test. **e-f**, Cell topology and eccentricity of st. 8 *Perlecan-depleted* follicles resembles WT. **g**, Cell orientation distribution in st. 8 *Perlecan-depleted* follicles resembles *fat2*. **h**, *10XSTAT-GAL4* drives transgene expression predominantly in follicle anterior at st. 7-8, where BM is soft and cell reorientation takes place. **i, j**, Depletion of *Rack1* in follicle anterior region is sufficient to induce elongation defects. Error bars in **d-g**, s.e.m. n, Sample size. NS, not significant, *P<0.5, **P<0.01, ***P<0.001, ****P<0.0001.

Finally, we investigated the relationship between Src activity, cellular orientation, and BM stiffness. Src is considered a mechanotransducer ^26^, and interestingly pSrc levels decrease as follicles develop, anticorrelating with increasing BM stiffness (**Fig. 7d**). Moreover, acute collagenase treatment caused a significant increase in pSrc (**Fig. 7d**), suggesting that cellular Src activity could be negatively regulated by BM stiffness. To test this proposition further, we analyzed elongation-defective follicles depleted of the ECM component Perlecan, which undergo rotation but display a uniformly soft BM (**Supplementary Fig. 3h**) ^19^. Indeed, *Perlecan*-depleted follicles also showed significant increases in pSrc levels (**Fig. 7d**), while increasing BM stiffness by overexpression of EHBP1^8,27^, which enhances Collagen IV fibril formation, had the opposite effect (**Fig. 7d**). Morphometric analysis showed that cell topology and eccentricity of *Perlecan*-depleted follicles resembled WT (**Fig. 7e-f**), but cell orientation at st. 8 was defective, displaying a distribution very similar to that seen in follicles depleted of *fat2* (**Fig. 7g**).

If BM stiffness regulates Src-mediated changes in cell orientation, then Rack1 should be required for follicle elongation primarily at the anterior, an area of soft BM that is the predominant site of reorientation. To test this, we created a GAL4 driver (10XSTAT-GAL4) that is preferentially expressed in anterior follicle cells at st. 7-8 (**Fig. 7h, Supplementary Fig. 4j**). Consistent with the model, RNAi-mediated depletion of Rack1 via this driver was sufficient to generate elongation defects, with a severity comparable to depletion throughout the follicle epithelium (**Fig. 7i-j**). Together, these data indicate that differences in the mechanical status of the BM, signaled via Src, regulate cell reorientation to allow tissue elongation.

## DISCUSSION

Mechanical regulation of animal tissues is a current frontier of biology, as new tools and model systems allow interrogation of unappreciated phenomena ^28-31^. An emergent model for morphogenesis shaped by extracellular as well as intracellular forces is the Drosophila follicle. Here we use *in toto* image analysis to define the cell dynamics that elongate this organ and to probe their regulation by mechanical properties of the extracellular matrix. We find that polarized reorientation of a subset of cells, involving the shift of their long axes from latitudinal towards AP meridional, is a major driver of elongation. This process fails when BM mechanics are compromised, and is regulated by the Src tyrosine kinase, which controls remodeling of cell-cell junctions. This unusual mechanism highlights differences between the elongation of bounded tissues and ‘edgeless’ acinar epithelia.

All three morphogenetic behaviors that are engines for tissue elongation in other systems –cell division, cell shape changes, and cell rearrangement- are seen during follicle elongation. Our manipulations show that oriented cell division is not essential for elongation, and its absence can be compensated by other polarized cell behaviors, as also seen in e.g. the pupal thorax ^32^. Additionally, an initial cell shape anisotropy --lengthening around the organ circumferential axis-- is not perturbed in a ‘round egg’ mutant, while tissue rotation alone is insufficient to direct elongation-driving cell dynamics.

Instead, our data suggest that the major tissue-elongating behavior involves changes in cell orientation. In particular, latitudinally-oriented cells in the follicle anterior shift the direction of their long axis to become more AP-oriented. Without changing average cell shape, this reorientation of cell anisotropy changes neighbor relationships, and results in net cellular intercalation along the AP axis. It is important to note that, in a 3D ellipsoid, large changes in overall aspect ratio along Cartesian axes can result from smaller changes in the ratio of meridian and equatorial arc lengths. Thus, more limited rearrangements may account for elongation.

The relative paucity of frank intercalation events seen in the follicle contrasts not only with the rapidly developing Drosophila germband, but also with other elongating tissues such as the Drosophila pupal wing and thorax and the vertebrate neural tube and mesoderm^33^. Three major differences bear consideration here. First, significant growth (~9-fold) of both epithelial cells and the follicle overall occur during the 24-34 hours ^11,34^ that span st. 4-8, while the other systems largely rearrange a constant tissue volume.

Second, the other systems have defined boundaries to impose vectorial orientation of cell behaviors, while the topologically continuous follicle epithelium lacks a boundary in the equatorial axis. Third, follicle elongation does not require conventional PCP morphogenetic signaling, nor is there evidence of PCP Myosin localization that remodels cell junctions ^8,20,35^ Instead, the instructive force seems to be patterned anisotropic resistance to growth ^8^, wherein softer BM at the poles triggers regional changes in cell behavior; indeed, the largest changes in follicle cell orientation are seen at the anterior pole, and genetic manipulation in this region alone can prevent elongation. These differences with bounded epithelia undergoing elongation via convergent extension emphasize the new perspectives required for analysis of elongating tubular organs.

How do cells sense BM stiffness to change their orientations? The elongation-defective phenotype of Rack1-depleted follicles points to one mechanism involving Src. Rack1 is a Src-inhibiting protein, and its loss, with associated increases in cellular Src activity, perturbs follicle elongation. In Rack1-depleted follicles, initial cell orientation is not greatly altered, but anterior follicle cells remain static and are unable to shift their orientation during stages 6-8. This suggests that proper regulation of Src is required for the dynamic reapportionment of cell shape that is reflected in this switch. One familiar regulatory target of Src is integrins and their associated proteins ^36,37^, but no defects in integrin-dependent cell migration are seen when Rack1 is depleted from follicle cells. However, Src has also been implicated in directly regulating AJ remodeling through effects on Ecad trafficking ^38-40^, and FRAP analysis reveals compromised Ecad dynamics when follicles cells lack Rack1. Src is considered a mechanotransducer, and pSrc levels anticorrelate with BM stiffness in WT, mutant and manipulated follicles. We therefore propose that Src mediates the instructive cue provided by BM stiffness, inducing AJ remodeling to drive morphogenesis.

The defective tissue topology and reduced Ecad mobility seen in Rack1-depleted follicles suggests that Src is required for both elongation-driving and tissue disorder-minimizing cell rearrangements. This role of Src in follicle elongation raises interesting parallels with a second ‘edgeless epithelium’: the Drosophila trachea. In this established tubulogenesis model, gain as well as loss of Src42 activity results in shortened but broader tubules, which result from inappropriately oriented cells ^41,42^. Interestingly, depletion of Src42 from the follicle, like its activation, also induced elongation defects (**Supplementary Fig. 4i**), while Src controls tracheal Ecad dynamics ^43^, perhaps in response to an ECM, albeit apically localized ^41,42,44^. Thus, in both organs Src42 could mediate stiffness-cued AJ dynamics that change the orientation of cell eccentricity. Since mammalian kidney tubule development also shows Src-dependence ^45^, these results suggest that ECM-mediated control of cell junctions via Src may be a general mechanism for morphogenesis of edgeless epithelia.

The data described in this work stem from a comprehensive analysis of follicle morphogenesis, which was enabled by ImSAnE analysis software. ImSAnE allowed identification of critical cell dynamics near the follicle poles, which are most subject to distortion from conventional analyses, and distinguished the cellular basis underlying overtly similar elongation phenotypes, i.e. between *Rack1* and *fat2* follicles. Nevertheless, some of the ImSAnE-based morphometric findings are concordant with those recently reported based on conventional imaging ^20,46,47^. This work focuses on st. 4-8, and does not address cell dynamics that elongate the follicle prior to and following these stages ^20,48^. However, it does provide a framework for true *in toto* analysis of this simple organ, at multiple developmental stages and genotypes. As genetic screens (this work, ^49^) and biophysical studies^8,19,50^ in the follicle are extended, the quantitative imaging platform reported here will provide a bridge towards mathematical and mechanical modeling of this flourishing system for *in toto* organ morphogenesis.

## Acknowledgments

Experiments using Zeiss Lightsheet Z.1 were conducted at the UC-Berkeley Molecular Imaging Center (RRID:SCR_0122850), the Cancer Research Laboratory, and the Helen Wills Neuroscience Institute (HWNI), with training and assistance from Holly Aaron and Jen-Yi Lee. We thank Brian Calvi, Shigeo Hayashi, Tetsuya Kojima, Greg Beitel, Yang Hong, TRiP at Harvard Medical School (NIH/NIGMS R01-GM084947), Kyoto Stock Center, and Bloomington Drosophila Stock Center (NIH P40OD018537) for providing fly stocks and reagents, and Laura Mathies for cloning 10XSTAT-GAL4 with advice from Ryan Boileau, Martin Zeidler, and Erika Bach. This work was supported by NIH RO1 grants GM068675 and GM111111 to D.B. and 4R00HD088708-03 to S.J.S

## Author Contributions

DYC and JC conducted the experiments; SJS developed analytical tools; DYC and DB designed the experiments and wrote the paper.

## METHODS

### Fly strains and husbandry

The following Drosophila strains were obtained from the stock center: *Jupiter-GFP* ^51^ (Flybase ID: FBst0006836), *mCherry-RNAi (control)* (Flybase ID: FBst0035785), *Fat2-RNAi* (Flybase ID: FBst0040888), *Rack1-RNAi* (Flybase ID: FBst0034694), *Perlecan-RNAi* (Flybase ID: FBst0029440), *Src42A-RNAi* (Flybase ID: FBst0044039), *Rack1^1.8^-FRT40A* ^52^, *UAS-Src42A-CA* (Flybase ID: FBst0006410), *P{tubP-GAL8Cf^s^}20* (Flybase ID: FBst0007019), *UAS-mcherry-Ehbp1* (Flybase ID: FBst0067145) ^53^, and *w1118* (Flybase ID: FBst0003605). UAS-fzr ^54^ is a gift from Brian Calvi; *Src42A^26-1^* is a gift from Greg Beitel ^23,42^; *UAS-Rack1.ORF* is from FLY-ORF # F001448; *Ecad-GFP* is a gift from Yang Hong ^25^; *GR1-GAL4, UAS-FLP* is a gift from Trudi Schüpbach; *Indy-GFP and vkg-GFP* are from Flytrap ^55^, *yw* is from Tom Neufeld, and *trafficjam-Gal4 (tj-GAL4)* from Kyoto Stock Center. 10XSTAT-GAL4 was created by synthesizing a 2250bp fragment containing five tandem copies of the Socs36E enhancer ^56^ and cloning into pAttB-Gal4. Detailed genotypes used in each figure are listed in **Supplementary Table 1**.

Adult flies were maintained at 25°C unless otherwise noted. Adult females were flipped onto fresh food daily for 1-2 days and were fed with yeast paste overnight before dissection.

*Fat2* follicles are RNAi-depleted, whose phenotype parallels molecularly characterized null alleles ^8^. *tj-GAL4, P{tubP-GAL8C^ts^}20; UAS-fzr (j^ts^UAS-fzr)* along with control follicles used in **Fig. 5e-f** were shifted to 29° for 17.5 hours before analysis. *tj^ts^* follicles used in **Supplementary Fig. 3i-j** were shifted to 29° for 25 hours before analysis. *tj^ts^>Rack1-RNAi* follicles used in **Supplementary Fig. 4f** were shifted to 29° for 24 hours before analysis. *10XSTAT-GAL4; P{tubP-GAL80^ts^}* follicles used in **Fig. 7h-j and Supplementary Fig. 4j** were shifted to 29° for 25 hours before analysis.

### Follicle staging

To objectively stage follicles, we compared morphological criteria ^5^ to our quantitative morphometric data on follicle volume and epithelial cell number (**Supplementary Fig. 1**). For stages with definitive morphological criteria, we generally found good accordance with our data, allowing consistent boundaries based on follicle cell number to mark the end of st. 4, 7, and 8. However, st. 5 and 6 are not clearly distinguished by morphology, and there is debate about whether cell proliferation ends prior to st. 6 or continues during this stage ^11,57,58^. We found that cell counts of follicles deemed st. 6 by the criteria of Jia et al. ^58^ contained well fewer than 900 cells. Moreover, the frequency of such follicles *in vivo* (our results and ^34^) is higher than would be expected for standard estimates of st. 6 duration ^11^. These standard estimates of developmental stages are based on transplant of a single germarium into the abdomen of an *ovoD1* female host, and are likely faster than follicles in an entire native ovary *in vivo* ^59^. Using 9.6 hours as the average length of a follicle cell cycle ^60^ the durations of st. 4, 5 and 6 based on our epithelial cell counts are 9.6, 4.8, and 8.4 hours respectively, in reasonable agreement with the durations suggested based on relative follicle representation in WT hosts ^34^. Due to the wide ranges in volume observed in classically defined st. 6 follicles, and the accelerated rate of elongation in larger follicles of this group, we divided them into substages 6A and 6B using cell number criteria. We also assessed convenient staging parameters in single confocal cross-sections when complete 3D follicles are not imaged. Like Dai et al. ^61^, we found that nurse cell nuclear diameter allowed reasonable staging, and our measurements are close to theirs. Jia et al. ^58^ used cross-sectional area of follicles, albeit with values discrepant with our data likely due to differences in mounting and imaging; we found that this parameter did not reliably distinguish follicles of different stages as determined by cell number and morphological criteria.

### Immunostaining and Imaging

Immunostaining and imaging were executed as previously described ^7,21^. Antibodies used are listed in **Supplementary Table 2**. *Ex vivo* follicle culture was performed as previously described ^8^, with osmolarity adjusted to 260 Osm/L. Fluorescent images were acquired on Zeiss LSM700 confocal microscope with LD C-Apochromat 40x/1.1 W Corr objective. Imaging for cell division orientation was performed using selective plane illumination microscope (SPIM) Zeiss LightSheet.Z1 with illumination lens 10X/0.2 and Lightsheet Z.1 detection optics 20X/1.0 W. Follicles were manually dissected and embedded in 0.5% low melting point agarose with complete media contained in a 0.6mm in diameter glass capillary (Brand GmbH). Mounted follicles were immersed into a chamber filled with media without insulin supplement.

### Morphometrics extraction

We used ImSAnE to unroll the follicle epithelium as previously described ^7,10^, with all measurements computed using the metric tensor. Follicle anterior-posterior (AP) orientation was defined either automatically by the longest axis or manually based on the position of the polar cells identified by anti-FasIII staining at the stages when follicles are close to spherical. Length along the AP meridian was computed by determining the anterior and posterior poles as extreme points along the AP axis, and then measuring the length of the geodesic connecting these two points. latitudinal length was computed by choosing an arbitrary point midway along the meridian, and measuring the length around the circumference passing through this point. Follicle aspect ratio is defined as the ratio of AP length to the length of future dorsoventral (DV) axis. Aspect ratios were computed from unflattened 3D stacks, except for **Fig. 7j** and **Supplementary Fig. 4f** which used single confocal sections of preparations with standard mounting ^8^. Follicle volume was computed by adding up all pixels contained within the 3D surface. The number of cells along the AP meridian and the equatorial circumference was determined by first segmenting all cells in a pullback. Cells whose centroids are a given distance away from the line used to measure the meridional and latitudinal length were then counted accordingly. For fusing 3D representations of the surface, pullbacks showing anterior, posterior, and two cylinder projections were first segmented, and then stitched together in 3D. The pullbacks were based on a maximum intensity projection of 3 x 0.5 μm thick pullbacks from 3-4 μm inward of the basal-most surface extracted. Cell segmentation used simple thresholding of prediction maps obtained by using Ilastik ^62^. The thresholded images were treated with standard morphological operations to obtain skeletonized cell outlines. Automated methods for detecting branching points then enabled construction of a lattice. The lattice consists of a lookuptable, with vertices, bonds connecting vertices, and cells arranging bonds (cell-cell interfaces) to a closed outline. The number of neighbors in each cell was determined by the number of bonds. Surface area was obtained by integrating the square root of the metric tensor over the cell. Summing all individual cell areas gave total follicle surface area. Cell eccentricity is determined based on the long and short axis of the cell, while cell orientation is defined as the angle between the long axis of the cell and the circumferential axis of the follicle.

### BM Stiffness Assays

Basement membrane stiffness was measured by Atomic Force Microscopy (AFM) as described previously ^8^.

### Fluorescence Recovery After Photobleaching (FRAP) analysis

FRAP was performed with Zeiss LSM 700 with LD C-Apochromat 40x/1.1 W Corr objective at room temperature (23.3°C). Pre- and post-bleaching images were captured every 1 sec for 5 min with a pixel resolution of 512×269 pixels (0.16 μm/pixel) and scan time 821.67 msec. For bleaching, an elliptical region of interest (ROI) ranging from 4.8-7.7 μm^2^ covering circumferentially oriented junctions was bleached twice by 488nm (10 mW) solid-state laser (70% laser power) with pixel dwell time 100.85 μsec. Both drifts in time lapse images and follicle rotation were corrected and reregistered with Fiji plugin StackReg followed by manual inspection and correction for each timeframe. Intensity of ROIs were measured using Fiji Mean gray value measurements. Normalized recovery curves were fitted by Prism with the following formula for one phase association:

Y0 +; (Plateau-Y0)*(1 - exp(-K*X))), where Y0 is the Y value immediately post bleaching, Plateau is the Y value at infinite times, K is the rate constant, half-time is ln(2)/K, mobile fraction is Plateau-Y0.

### Fluorescence intensity and follicle velocity measurements

Fiji ^63^ was used to measure the intensity of apical Src and phosphorylated Src (pSrc) intensity. A 10-pixel thick line along the apical membranes was drawn with freehand line tool, followed by mean gray value measurements. Intensities were normalized based on the mean intensity of control st. 8 follicles from the same batches under the same imaging settings.

Follicle rotation velocities were measured by the circumferential displacement of plasma membranes visualized with Indy-GFP at the equator.

### Collagenase treatment

*ex vivo* cultured follicles were treated with collagenase as previously described^8^, followed by 3x washes with Schneider’s media and 35min of incubation at 25°C before fixation.

### Data presentation

Opacity projection was generated by Volocity 5 (PerkinElmer). ImSAnE surface 3D projection was generated by plotting cylinder projection onto fitted 3D structure. Max Z projection was generated by Fiji Z projection-Max intensity function. Data were analyzed and charts displayed using MATLAB 2015a and 2018a (Mathworks), Excel (Microsoft), Prism 6 (Graphpad) and ggplot2 ^64^ in RStudio ^65^. Figures were assembled with Adobe Illustrator CC 2018 (Adobe).

### Statistics and reproducibility

Neither sample size estimate, randomization of samples or blinding was performed. Data normality and variation estimation were not performed. Unless noted, error bars in charts represents s.e.m. Statistical significance assembled was assessed by two-sided Welch’s unequal variances *t*-test, while two-sample Kolmogorov-Smirnov test was used to analyze estimated probability distribution functions in cell eccentricity and cell orientation. All experiments were replicated with distinct biological samples at least three times. Data were only excluded when samples are damaged or the time-lapses showed significant drifts during imaging.

### Code availability

Customized computer codes used in this study are available upon request.

